# Autophagy-dependent alternative splicing event produces a more stable ribosomal protein S24 isoform that aids in hypoxic cell survival

**DOI:** 10.1101/2023.09.25.559338

**Authors:** Jenna Kerry, Erin J. Specker, Morgan Mizzoni, Andrea Brumwell, Leslie Fell, Jenna Goodbrand, Michael N. Rosen, James Uniacke

**Affiliations:** Department of Molecular and Cellular Biology, University of Guelph, Guelph, Ontario, Canada

**Author notes:** Corresponding author: Department of Molecular and Cellular Biology, University of Guelph, 50 Stone Road East, Guelph, Ontario, Canada N1G 2W1.

**Keywords:** Ribosome, Hypoxia, Splicing, Autophagy, Stress

## Abstract

Overlapping or convergent stress-activated molecular pathways exist to coordinate cell fate in response to stimuli such as hypoxia, oxidative stress, DNA damage, and unfolded proteins. Cells can remodel the splicing and translation machineries to mount a specialized gene expression response to certain stresses. Here, we show that hypoxic human cells in 2D and 3D culture models increase the relative abundance by 1.7- to 2.6-fold and 4.7- to 11.5-fold, respectively, of a longer mRNA variant of ribosomal protein S24 (RPS24L) compared to a shorter mRNA variant (RPS24S) by favoring the inclusion of a 22 bp cassette exon. Mechanistically, RPS24L and RPS24S are induced and repressed, respectively, by distinct parallel pathways in hypoxia: RPS24L is induced in an autophagy-dependent manner, while RPS24S is reduced by mTORC1 repression and in a HIF-dependent manner. RPS24L is a more stable mRNA in hypoxia and produces a more stable protein isoform compared to RPS24S. Cells overexpressing RPS24L display improved survival and growth in hypoxia relative to control cells and cells overexpressing RPS24S, which display impaired survival. Previous work from our group showed a correlation between RPS24L levels and tumor hypoxia in prostate cancer. These data highlight RPS24L as a stress-induced alternative splicing event that favors hypoxic cell survival, which could be exploited by cancer cells in the tumor microenvironment.

## INTRODUCTION

Cells routinely encounter various stresses that require either an adaptive response to return to homeostasis or a decision to trigger cell death. Overlapping or convergent stress-activated molecular pathways exist to coordinate cell fate in response to stimuli such as hypoxia, oxidative stress, DNA damage, and unfolded proteins. One such pathway is mediated by the nutrient and energy sensor mammalian target of rapamycin complex 1 (mTORC1) that receives inputs from several different sources to regulate cell growth (1). The heat shock and unfolded protein responses are well-characterized pathways that rely on increased chaperone activity to enhance the protein folding capacity of the cell. Autophagy is another system that is activated upon stress exposure and mTORC1 inhibition to remove or recycle damaged cellular components. If the protective or survival pathways cannot overcome persistent irresolvable stress, then different forms of cell death will be activated.

Hypoxia, or low oxygen availability, is an energy-limiting stress that represses mTORC1 and induces autophagy, among many other signaling cascades (2,3). Central to the hypoxic gene expression response are the hypoxia inducible factors (HIF) 1 and 2, heterodimeric transcription factors with α and β subunits that share most of their transcription targets but bind to distinct targets too. HIF-2α duals as a translation initiation factor in non-canonical cap-dependent translation in partnership with eIF4E2 (4,5). Indeed, mTORC1 repression in hypoxia shifts the translational apparatus away from canonical eIF4E-mediated cap-dependent translation toward cap-independent translation and the aforementioned eIF4E2-mediated pathway (4,6,7). Hypoxia drives many physiological and pathophysiological process and is a major characteristic of the tumor microenvironment that drives cancer progression (8).

At the heart of all protein production are macromolecular complexes called ribosomes that are made of ribosomal proteins (RP), ribosomal RNA (rRNA), and many non-ribosomal proteins. In mature ribosome, rRNAs act as the catalytic core, whereas RPs facilitate the processing and folding of precursor rRNA into mature rRNA via endonucleolytic and exonucleolytic cleavages (9). Historically, the ribosomal subunits themselves were thought to only have a constitutive role in translation and were viewed as passively active proteins that non-specifically translate mRNA due to their vital role in protein synthesis among all kingdoms of life (10). However, rRNA and RPs can be subjected to modifications that possibly play a role in translation regulation (11). Recent evidence suggests that eukaryotic ribosomes are not merely passive static complexes but can be heterogenous and selectively translate specific mRNAs in response to various stimuli (12).

Our group previously screened 68 alternative splicing events (ASEs) in RP transcripts and found that only one was altered by hypoxia that also affected the protein coding region (13). This ASE in RPS24 increased the retention of a 22 bp cassette exon by 1.4- to 2.1-fold in hypoxic cells and up to 10-fold in spheroids (*in vitro* tumor models) of four different cell lines relative to normoxic cells. This longer RPS24 mRNA variant (RPS24L) contains a premature stop codon that shortens the encoded protein by three amino acids at the C-terminus compared to the protein produced by the short RPS24 mRNA variant (RPS24S). We showed that both RPS24L and RPS24S protein isoforms incorporate into actively translating ribosomes (13), presumably either one or the other. Alternative splicing has been proposed as the 11th hallmark of cancer (14) and we showed that RPS24L levels correlate with hypoxia in human prostate tumor samples (13). Without reference to splicing isoforms or hypoxia, other groups have found that global RPS24 levels are high in prostate (15) and hepatocellular cancer (16), and show that RPS24 is an oncoprotein (17). Therefore, we set out to investigate the stimuli and mechanisms that contribute to the increase of RPS24L relative abundance compared to RPS24S and the biological impact this has on hypoxic cells.

Here we show that spheroid cells increased the relative abundance of RPS24L gradually over a time course across four different cell lines. This relative abundance was increased by other spheroid and tumor microenvironment stimuli such as acidosis, apoptosis induction, and chronic hypoxia. Mechanistically, RPS24L and RPS24S were induced and repressed, respectively, by distinct parallel pathways in hypoxia: RPS24L was induced in an autophagy-dependent manner, while RPS24S was reduced in a HIF-dependent manner and during mTORC1 repression. RPS24L was a more stable mRNA in hypoxia that produced a more stable protein isoform. We show that cells overexpressing RPS24L displayed improved survival and growth in hypoxia relative to control cells and cells overexpressing RPS24S, which displayed impaired survival. These data highlight RPS24L as an alternative splicing event that favors hypoxic cell survival, which could be exploited by cancer cells in the tumor microenvironment.

## RESULTS

### The formation of cells into spheroids increases the relative abundance of the RPS24 long mRNA variant compared to the short

We previously showed that hypoxia induces RPS24L by 1.4- to 2.1-fold in cell monolayers and by 4- to 10-fold in five-day old spheroids of HEK293 cells and cancer cell lines U87MG glioblastoma, HCT116 colorectal carcinoma, and PC3 prostate cancer relative to normoxic monolayers (13). We generated spheroids from these four cell lines and monitored the abundance of RPS24L and RPS24S across seven days of growth. Spheroids generate an internal hypoxic microenvironment that increases over the course of their growth; therefore, they do not require hypoxic incubation (18). After one day, spheroids displayed a 3- to 6-fold significant increase in RPS24L compared to monolayers in all four cell lines (Fig. 1A-D). With gradual increases over time, at the final day of measurement (seven days), spheroids displayed a 6- to 12-fold significant increase in RPS24L compared to monolayers in all four cell lines (Fig. 1A-D). In these seven-day old spheroids, RPS24S was reduced relative to monolayers in some cell lines such as U87MG and PC3 (Fig. 1B and D). Therefore, RPS24L induction and in some cell lines RPS24S repression contributed to the rising relative abundance between these mRNA variants, which increased at a higher rate between days five and seven than between any earlier time points in all four cell lines (Fig. 1A-D).

**Figure 1.**
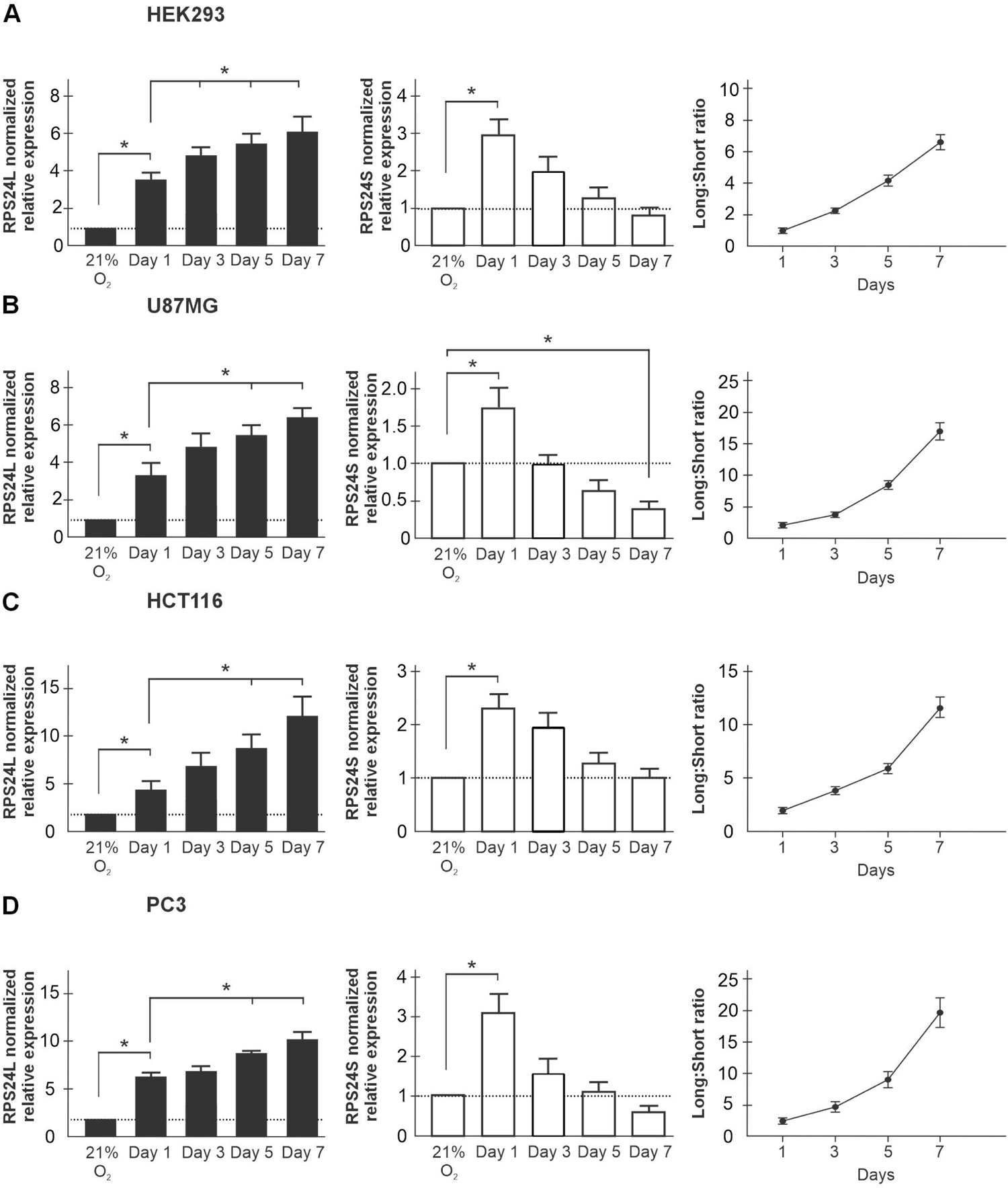
The formation of cells into spheroids increases the relative abundance of the RPS24 long mRNA variant compared to the short. Spheroids from (A) HEK293, (B) U87MG, (C) HCT116, and (D) PC3 were lysed on days 1, 3, 5, and 7 post-seeding. The expression of the long and short RPS24 mRNA variants was measured using qRT-PCR and variant-specific primers. The ΔΔCt method was used, normalizing to reference genes *RPLP0* and *RPL13A*, and the normalized expression made relative to each variant in a normoxic monolayer. (n ≥ 4), mean normalized relative expression ± s.e.m. Normalized relative expression was used to calculate the ratio of long to short variants. One-way ANOVA with Tukey’s HSD test was performed on the ΔΔCt values. * = P < 0.05.

We noticed that RPS24S levels significantly increased, albeit to a lesser extent than RPS24L, by 1.5- to 3-fold in one-day old spheroids in all four cell lines (Fig. 1). This was not captured in our previous report where RPS24 variant levels were measured only after five days of spheroid formation (13). Furthermore, both RPS24 variants increased by 1.5- to 2-fold one day after seeding whether lifted with trypsin or scraped in phosphate buffered saline (Fig. S1A-B). Including the seeding (24 h) and hypoxic incubation (24 h), it is important to note that both normoxic and hypoxic monolayer controls used in our previous and current study to measure RPS24 variant levels are 48 h after seeding. Therefore, it appears that there is a plating-induced stress that induced both RPS24L and S variants within 24 h of seeding into monolayers or spheroids (Fig. S1A-B). RPS24S levels returned to baseline or lower after three to five days of spheroid formation, while RPS24L continued to increase, which is the main focus of this study.

In Figure 1, each RPS24 mRNA variant was normalized to their respective normoxic monolayer. When we compared the relative abundance of each variant within normoxic monolayers we noticed that all four cell lines had substantially different RPS24L to S baseline ratios: 1.25 ± 0.05 in HEK293, 0.44 ± 0.04 in U87MG, 0.24 ± 0.01 in HCT116, and 3.6 ± 0.7 in PC3. U87MG and HCT116 had more RPS24S in normoxia, PC3 had more RPS24L, and HEK293 had approximately equal proportions of both. However, once exposed to hypoxia for 24 h, the RPS24L/RPS24S ratios all increased by 1.7- to 2.6-fold regardless of the baseline normoxic ratio (Table 1). Furthermore, formation into spheroids after seven days produced increases to the RPS24L/RPS24S ratio from 4.7- to 11.5-fold in all four cell lines. These data suggest that cells of different origin have a unique RPS24L/RPS24S ratio, but hypoxia and the stimuli within the spheroid microenvironment increase this ratio in a consistent manner.

**Table 1.** RPS24 long:short mRNA variant ratio increases consistently across four cell lines. The expression of the long and short RPS24 mRNA variants was compared within conditions of normoxic and hypoxic monolayers, and five-day old spheroids using qRT-PCR and variant-specific primers. N=3 shown as mean ± standard error. Fold increase of the RPS24 long:short mRNA variant ratio is shown relative to the normoxic monolayer.

| Cell line | Normoxia | Hypoxia | Spheroid |
| --- | --- | --- | --- |
| HEK293 | 1.25 ± 0.05 | 2.13 ± 0.12<br>(1.7-fold) | 7.35 ± 0.35<br>(4.7-fold) |
| U87MG | 0.44 ± 0.04 | 1.15 ± 0.10<br>(2.6-fold) | 7.48 ± 0.65<br>(17.0-fold) |
| HCT116 | 0.24 ± 0.01 | 0.39 ± 0.03<br>(1.7-fold) | 1.66 ± 0.23<br>(6.9-fold) |
| PC3 | 3.60 ± 0.67 | 9.39 ± 2.07<br>(2.6-fold) | 24.06 ± 0.77<br>(6.7-fold) |

### The RPS24 long to short mRNA variant ratio increases in low pH, chronic hypoxia, and staurosporine and can be reversed after spheroid disassembly

Spheroids increase the RPS24L/RPS24S ratio much more than in hypoxic monolayers (6- to 12-fold compared to 1.4- to 2.1-fold) relative to normoxic monolayers. We aimed to identify the stimuli within the spheroid microenvironment responsible for this larger increase by treating U87MG monolayers with several individual stimuli since their spheroids displayed the largest increase in RPS24L/S ratio. Due to the high glycolytic activity of tumor cells, low pH (5.6 to 6.8) is a hallmark within solid tumors (19). Only the lowest pH (5.8) significantly increased RPS24L by 1.6-fold relative to neutral pH 7.0 in normoxia (Fig. 2A). Of note, hypoxia and pH 5.8 alone caused similar increases to RPS24L, but a synergy was not observed. While lactic acid accumulates during glycolysis, it also acts as a signaling molecule (20). We treated normoxic and hypoxic cells with either lactic acid or sodium lactate and observed no change to RPS24L or RPS24S mRNA abundance (Fig. 2B-C). We next tested whether the increase in cell density in spheroids compared to monolayers was a signal that raised the RPS24L/S ratio. Cells were seeded as monolayers at equal density and allowed to grow for 1, 2, 3, 4, and 5 days (Fig. 2E). As monolayers increased in density, by days 4 and 5 both RPS24L and S significantly decreased in abundance relative to day 1, possibly due to signals to decrease proliferation and ribosome biogenesis (Fig. 2F). Incubating cells in hypoxia for seven days significantly increased RPS24L mRNA abundance by 1.7-fold relative to control “Day 1” normoxic cells, but this induction was not any higher than what we observed after 24 h of hypoxia (13). Cell density was equalized between Day 1 and Day 7 cells by performing seven two-fold serial dilutions and confirming visually before lysing. Spheroids undergo an early period (≤ 36-48 h from seeding) of apoptotic activity followed by a shift to increased decomposition and necrosis (21). The inducer of apoptosis staurosporine did not induce RPS24L in normoxic monolayers, but it did repress RPS24S (Fig. 2H). Therefore, apoptosis in the early period of spheroid formation is likely not responsible for RPS24L induction but could contribute to an increased RPS24L/RPS24S ratio. To test whether an excreted signal from spheroid cells contributes to RPS24L induction, we treated normoxic monolayers with media from five-day old spheroids. No significant changes to RPS24L or S were observed (Fig. 2I). We next disrupted calcium-dependent cell-cell contacts (i.e., cadherins) in spheroids with the chelating agent EGTA to test whether RPS24L induction could be reversed. Indeed, spheroids treated with EGTA displayed a significant decrease in RPS24L mRNA abundance almost back to baseline levels observed in normoxic monolayers (Fig. 2J). Even though the spheroid microenvironment as a whole is disrupted by EGTA and not only cell-cell interactions, these data show that RPS24L induction is not a one-way response and can be reversed. Further, RPS24L induction can occur in low pH and chronic hypoxia (168 h), but this was to the same extent as 24 h of hypoxic incubation and still much lower than what is observed in spheroids.

**Figure 2.**
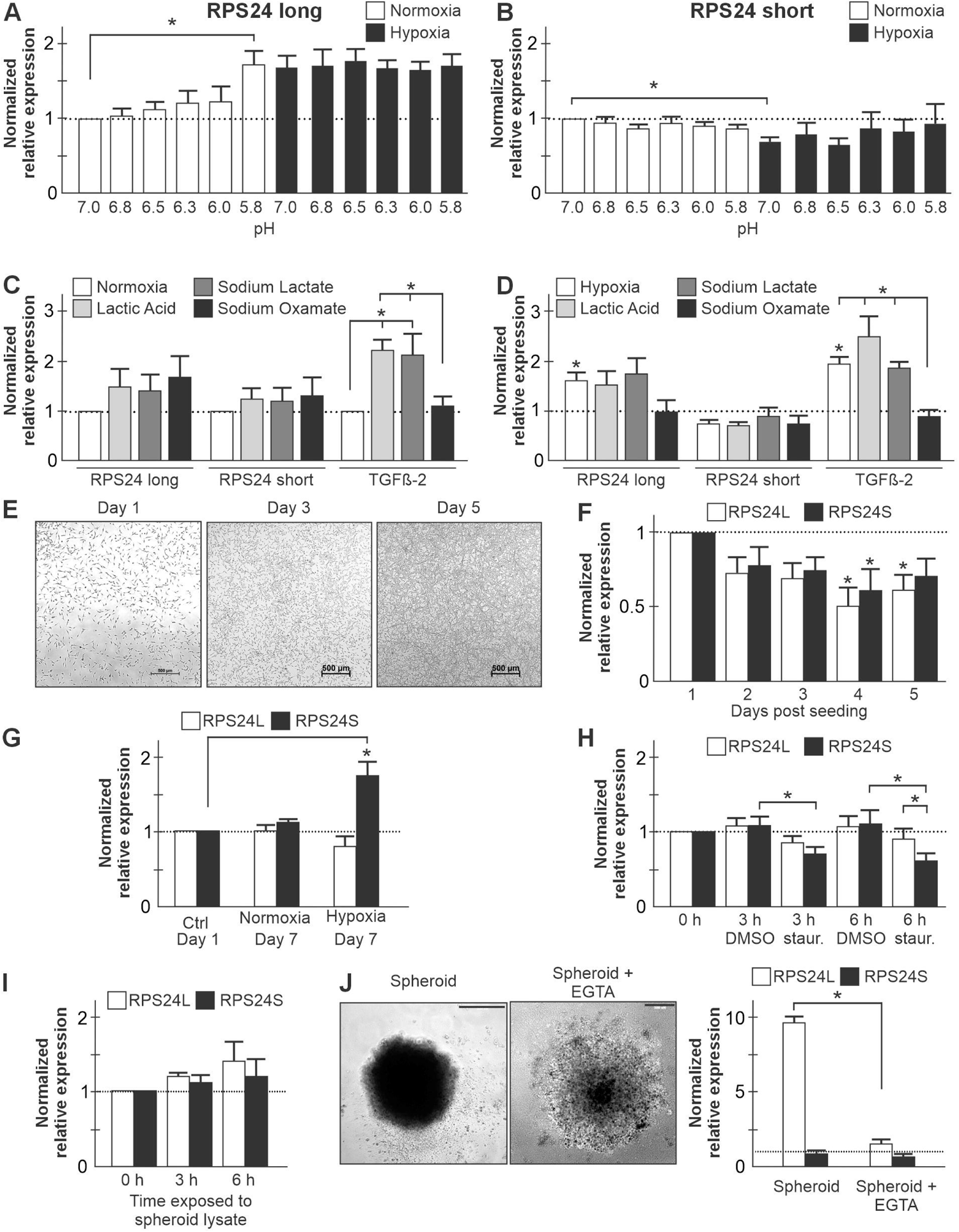
The RPS24 long variant is induced by low pH and chronic hypoxia, and its induction is reversed after spheroid disassembly. RPS24 long (A) and short (B) mRNA variant abundance was measured in normoxic and hypoxic cells treated for 24 h with acidosis-permissive media to produce pH ranges from 5.8 to 7.0 relative to normoxic cells at pH 7.0. Serum-starved normoxic (C) and hypoxic (D) cells were treated for 24 h with lactic acid, sodium lactate, and the negative control sodium oxamate. The mRNA abundance of the hypoxia- and lactate-responsive gene *TGFβ-2* and both RPS24 mRNA variants was measured and made relative to the normoxic control. (E) Representative images of cells on days 1, 3, and 5 of growth after an equal number of cells were seeded on the same day. (F) Cells were seeded from a serial dilution and lysed at one day intervals for five days. mRNA abundance of RPS24 long and short variants was measured relative to their day 1 value. (G) RPS24 long and short mRNA variant abundance was measured in cells incubated in hypoxia and normoxia for seven days relative to control (Ctrl) cells considered Day 1 cells (24 h seeding + 24 h normoxic incubation). (H) The mRNA abundance of RPS24 long and short variants was measured in cells treated with the inducer of apoptosis, staurosporine, and vehicle control DMSO for 3 and 6 h relative to the untreated control. (I) RPS24 long and short variant mRNA abundance was measured in cells treated for 3 and 6 h with media from five-day old spheroids after they were disassociated, and cells removed relative to untreated cells. (J) Cell-cell interactions were impaired with EGTA treatment in five-day old spheroids followed by measurement of RPS24 long and short variant mRNA abundance relative to normoxic monolayers. qRT-PCR data (n ≥ 3) displayed as mean normalized relative expression ± s.e.m. One-way ANOVA with Tukey’s HSD test was performed on the ΔΔCt values. * = P < 0.05. Scale bars, 500 µm. All experiments performed in U87MG glioblastoma cells.

### RPS24S repression, but not RPS24L induction, is HIF-dependent

Since hypoxia is a common feature of the tumor/spheroid microenvironment, we set out to investigate the mechanism used to repress RPS24S and induce RPS24L by chemically stabilizing or inhibiting the HIFs. Normoxic cells were treated for 24 h and 48 h with the HIF stabilizer Dimethyloxalylglycine (DMOG), an inhibitor of the prolyl hydroxylase domain-containing proteins. HIF activity was confirmed by the increase in GLUT1 mRNA, which is synthesized from a HIF-dependent promoter (Fig. 3A). Similar to what is observed in hypoxia (13) and spheroids (Fig. 1), DMOG treatment significantly reduced RPS24S mRNA abundance at 24 h and 48 h compared to vehicle control (Fig. 3A). However, RPS24L was only significantly induced (2.3-fold) after 48 h of DMOG treatment (Fig. 3A). Interestingly, when DMOG-treated cells were treated with HIF inhibitors, RPS24S repression was alleviated but RPS24L induction was not reversed (Fig. 3A). Similar results were observed in five-day old spheroids where the high relative level of RPS24L observed in Fig. 1B was not reduced by HIF inhibitors, but the repression of RPS24S mRNA abundance was significantly alleviated by more than 2-fold (Fig. 3B). Moreover, exogenous stable HIF-α subunits repressed RPS24S mRNA abundance in normoxia but had no effect on RPS24L (Fig. 3C-D). These exogenous HIF-α subunits contain proline to alanine mutations that stabilize them in normoxia by escaping proteasomal targeting via prolyl hydroxylases. These data suggest that hypoxic repression of the RPS24S mRNA variant is HIF-dependent, but the hypoxic induction of RPS24L is not.

**Figure 3.**
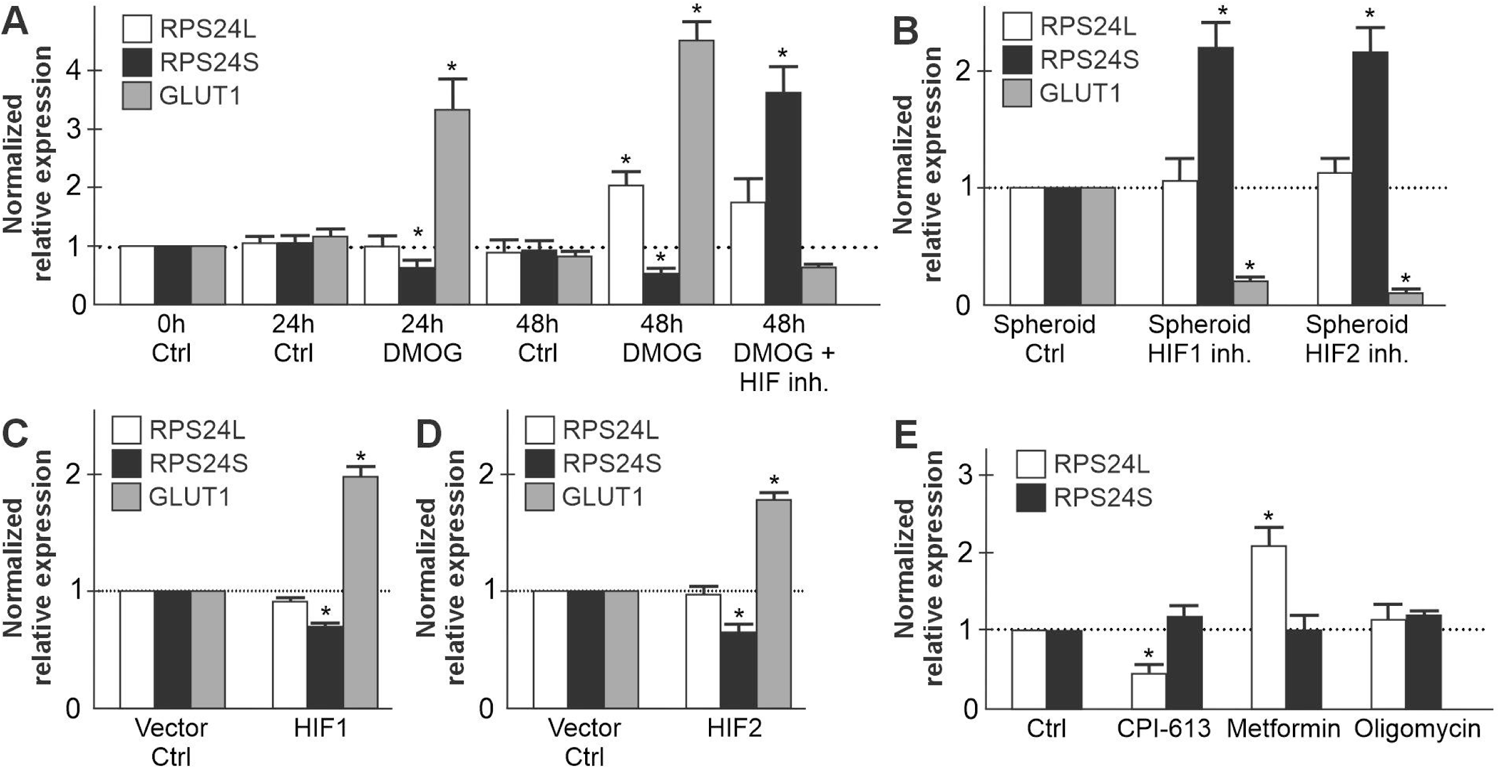
RPS24S repression, but not RPS24L induction, is HIF-dependent. (A) RPS24 long and short mRNA variant and GLUT1 mRNA (HIF-dependent control) abundance was measured in normoxic cells treated with the HIF stabilizer Dimethyloxalylglycine (DMOG), an inhibitor of the prolyl hydroxylase domain-containing proteins, for 24 and 48 h and made relative to untreated control (0 h Ctrl). Cells were also treated with a combination of DMOG and a cocktail of HIF-1 (Echinomycin) and HIF-2 (TC-S 7009) inhibitors. Statistical tests compare treated groups with vehicle only (DMSO) at the same time point. (B) RPS24L and RPS24S mRNA variant and GLUT1 mRNA abundance was measured in five-day old spheroids treated with either Echinomycin or TC-S 7009. Each transcript and statistical test were relative to spheroids treated with vehicle control (ctrl; DMSO). Normoxic cells were transfected with exogenous HIF-1α (C), HIF-2α (D), or their respective empty vector backbone control (Ctrl), and RPS24L and RPS24S mRNA variant and GLUT1 mRNA abundance were measured and made relative to Ctrl. Exogenous HA-HIF-α or FLAG-GFP-HIF-2α contain proline to alanine mutations that stabilize them in normoxia by escaping proteasomal targeting via prolyl hydroxylases. (E) RPS24 long and short mRNA variant abundance was measured in normoxic cells treated for 2 h with inhibitors of mitochondrial respiration with different modes of action: CPI-613, Metformin, or Oligomycin relative to vehicle control only treatment (Ctrl; DMSO). qRT-PCR data (n ≥ 3) displayed as mean normalized relative expression ± s.e.m. One-way ANOVA with Tukey’s HSD test was performed on the ΔΔCt values. * = P < 0.05. All experiments performed in U87MG glioblastoma cells.

The prolyl hydroxylases that stabilize the HIFs require α-ketoglutarate (α-KG) as a co-factor for activity. DMOG binds to and inhibits these enzymes by acting as an α-KG analog. However, because of this property, DMOG also inhibits α-KG dehydrogenase (α-KGDH) and mitochondrial respiration (22). We investigated whether the induction of RPS24L mRNA abundance after 48 h of DMOG treatment was due to the inhibition of mitochondrial respiration by treating normoxic cells with three different inhibitors: CPI-613 (specific inhibitor of α-KGDH), Metformin (mitochondrial complex I inhibitor), and Oligomycin (ATP synthase inhibitor). Inhibiting α-KGDH did not induce RPS24L, but instead reduced it (Fig. 3C). Oligomycin had no effect on RPS24L levels, but Metformin induced them by 2.1-fold relative to vehicle control (Fig. 3C). RPS24S levels were not affected by these inhibitors of mitochondrial respiration. These data suggest that the DMOG-dependent induction of RPS24L mRNA is not due to inhibition of α-KGDH or mitochondrial respiration in general, but perhaps due to some common effect shared between DMOG and Metformin.

### Hypoxic induction of RPS24L is dependent on autophagy, while RPS24S is repressed by mTOR inhibition

A shared property of DMOG and Metformin is their cytotoxic induction of autophagy (23,24). Cells treated with DMOG for 24 and 48 h potently stabilized the HIF-α subunits and displayed reduced p62 (Fig. 4A), its clearance an indicator of more active autophagic flux due to its presence in autophagosomes (25). Even lower p62 levels suggest higher autophagic flux at 48 h of hypoxia (Figure 4A). Due to the temporal expression of the HIF-α subunits and peak expression at ≤ 2 h (for HIF-1α) and ≤ 24 h (for HIF-2α), the hypoxic treatment did not capture peak HIF-α expression (26). These data convinced us to more thoroughly examine the link between autophagy and the induction of RPS24L to RPS24S relative abundance.

**Figure 4.**
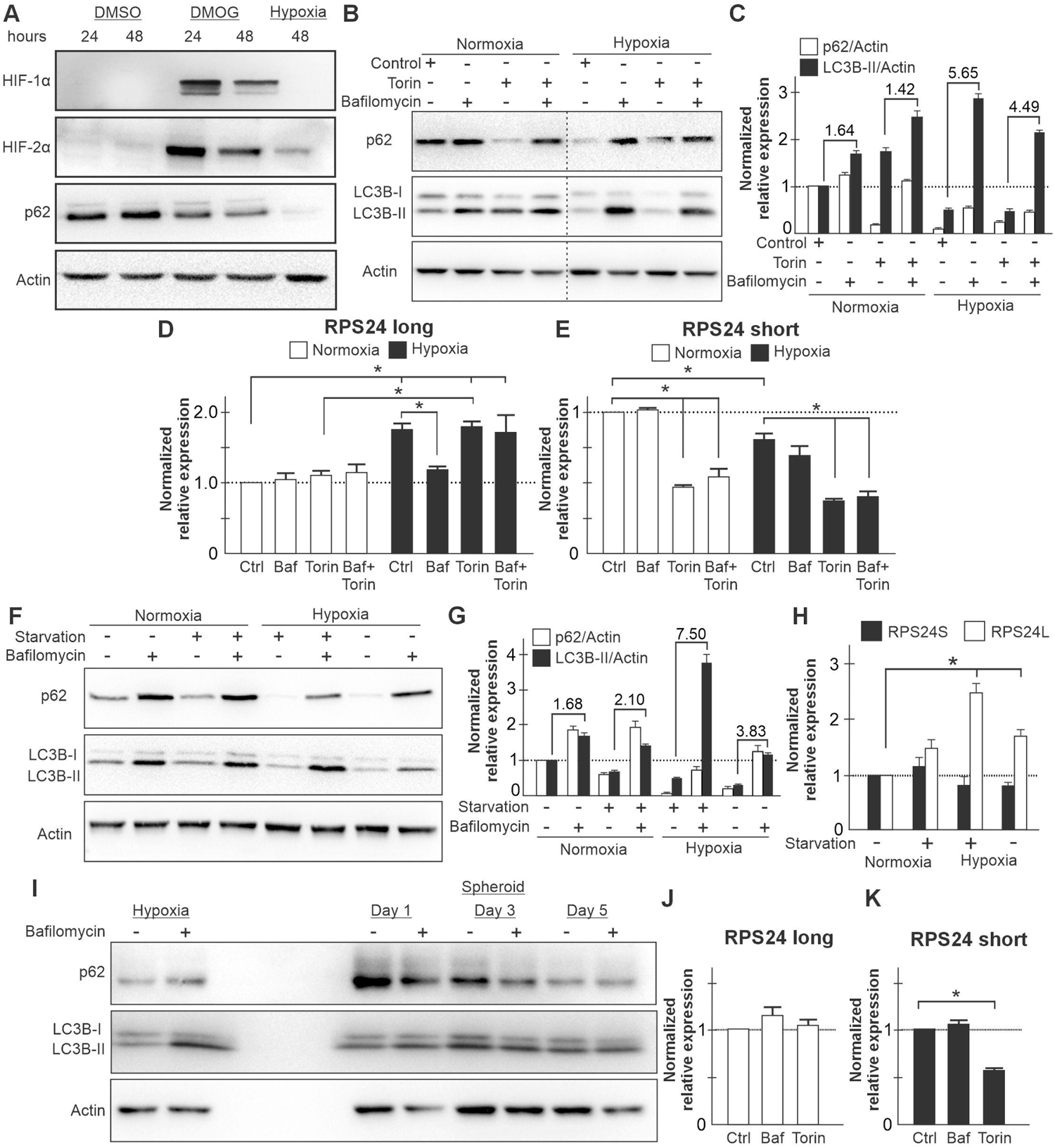
Hypoxic induction of RPS24L mRNA is dependent on autophagy. (A) Normoxic cells were treated with DMOG, or vehicle control (DMSO), for 24 and 48 h and compared to cells incubated in hypoxia for 48 h. Western blot used to detect HIF-1α, HIF-2α, and p62 (autophagy marker). (B) Normoxic and hypoxic cells were treated with Torin (inducer of autophagy), Bafilomycin (autophagy inhibitor), and/or vehicle control (DMSO) for 24 h. Western blot used to detect markers of autophagic flux (high activity → low p62, high LC3B-II +/− Bafilomycin ratio). (C) Quantification by densitometry of Western blots (n=3) in (B) using ImageJ software displayed as mean ± s.e.m. Ratio of LC3B-II levels +/− Bafilomycin is displayed. RPS24 long (D) and short (E) mRNA variant abundance was measured in normoxic and hypoxic cells treated with Torin, Bafilomycin, and/or vehicle control (Ctrl). (F) Markers of autophagy measured by western blot in normoxic and hypoxic cells without (-) or with (+) serum starvation and treated with or without Bafilomycin for 24 h. (G) Quantification by densitometry of Western blots (n=3) in (F) using ImageJ software displayed as mean ± s.e.m. (H) RPS24 long and short mRNA variant abundance was measured in normoxic and hypoxic cells without (-) or with (+) serum starvation. (I) Autophagy markers were detected in one-, three-, and five-day old spheroids with and without Bafilomycin treatment and compared to monolayers cultured in hypoxia for 24 h. RPS24 long (J) and short (K) mRNA variant abundance was measured in five-day old spheroids treated with Bafilomycin, Torin, and vehicle control (DMSO) for 24 h. qRT-PCR data (n ≥ 3) displayed as mean normalized relative expression ± s.e.m. One-way ANOVA with Tukey’s HSD test was performed on the ΔΔCt values. * = P < 0.05. All experiments performed in U87MG glioblastoma cells.

Autophagic flux was assessed in normoxic and hypoxic cells by measuring LC3B-II turnover. In the presence of a lysosomal degradation inhibitor such as Bafilomycin, LC3B-II accumulation can be an indicator of autophagic flux when compared to its baseline levels in the absence of Bafilomycin (25). The mTORC1 inhibitor Torin was used as an inducer of autophagy (27), while p62 levels were used as an alternate measure of autophagic flux. Hypoxia displayed a much higher autophagic flux (LC3B-II with/without Bafilomycin ratio of 5.65) compared to normoxia that had a ratio of 1.64 (Fig. 4B-C). Torin did not affect LC3B-II turnover but significantly reduced p62 levels in normoxia, which were rescued by Bafilomycin. Further, Torin did not appear to induce autophagy any more than hypoxia alone. We next examined the abundance of RPS24L and S in these same conditions. Bafilomycin prevented hypoxic cells from inducing RPS24L (Fig. 4D), but did not prevent the reduction of RPS24S (Fig. 4E). Torin had no effect on RPS24L levels (fig. 4D), but repressed RPS24S by over 2-fold independent of oxygen levels and autophagy (Fig. 4E). Bafilomycin did not prevent the hypoxic induction of RPS24L in the presence of Torin, although when comparing the Bafilomycin + Torin RPS24L mRNA abundance between normoxia and hypoxia there was not a significant hypoxic induction of RPS24L (Fig. 4D).

We next examined whether RPS24L could be induced by serum starvation, a potent inducer of autophagy (28). Indeed, serum starvation mildly induced autophagy in normoxia (2.10 compared to 1.68 LC3B-II ratio with/without Bafilomycin) and potently induced autophagy in hypoxia (7.50 vs. 3.83) (Fig. 4F-G). Serum starvation also reduced p62 levels in both normoxia and hypoxia (Fig. 4F-G). Normoxic serum starvation induced RPS24L by 1.5-fold ± 0.1 albeit not statistically significant (Fig. 4H). However, hypoxic serum starvation increased RPS24L mRNA abundance 2.52-fold ± 0.18 relative to normoxic cells in complete media, which is an additive effect above the hypoxic baseline RPS24L levels. Next, we sought to examine whether high levels of autophagy could be observed in spheroids, an environment that produced the highest observed RPS24L increase and RPS24L/S ratio relative to normoxic cells (Fig. 1). Unfortunately, autophagic flux could not be determined because Bafilomycin treatment did not produce LC3B-II accumulation in spheroids (Fig. 4I). This could be due to the density of spheroids since a slight accumulation of LC3B-II could be observed in one day old spheroids, which are looser. Since Bafilomycin did not appear to function in spheroids, it was not surprising that Bafilomycin did not reduce RPS24L mRNA abundance in five-day old spheroids (Fig. 4J). However, there was a gradual reduction in p62 across one-, three-, and five-day old spheroids suggesting autophagy is increasing as spheroids grow and could still likely be a factor in RPS24L induction considering the data presented in monolayers (Fig. 4D). Torin did produce a similar decrease in RPS24S mRNA abundance in five-day old spheroids (Fig. 4K) as observed in monolayers (Fig. 4E). Hypoxia is a known inducer of autophagy and inhibitor of mTOR (3,29). These data suggest that the hypoxic increase of RPS24L is dependent on autophagy but independent of mTOR, and RPS24S is reduced under mTOR repression independent of autophagy.

### The RPS24 long mRNA variant and the encoded protein isoform are more stable relative to the short isoform in hypoxia

While it is important to understand what environments and mechanisms are involved in promoting the inclusion of the 22 bp cassette exon in RPS24, we next examined if any cellular benefit could be observed. A transcript stability assay was performed in the presence of Actinomycin D using variant-specific primers in normoxic and hypoxic cells. Ribosomal protein mRNAs and proteins can have very long half-lives (30), and this is evidenced by their inclusion as unchanging reference genes in various experimental conditions in qRT-PCR assays. Both RPS24S and RPS24L mRNA variants were similarly stable in normoxia, displaying reductions of two-fold or less after 48 h of Actinomycin D treatment (Fig. 5A). However, in hypoxia, RPS24S was significantly less stable than RPS24L by two-fold or more at both 24 h and 48 h of Actinomycin treatment (Fig. 5B).

**Figure 5.**
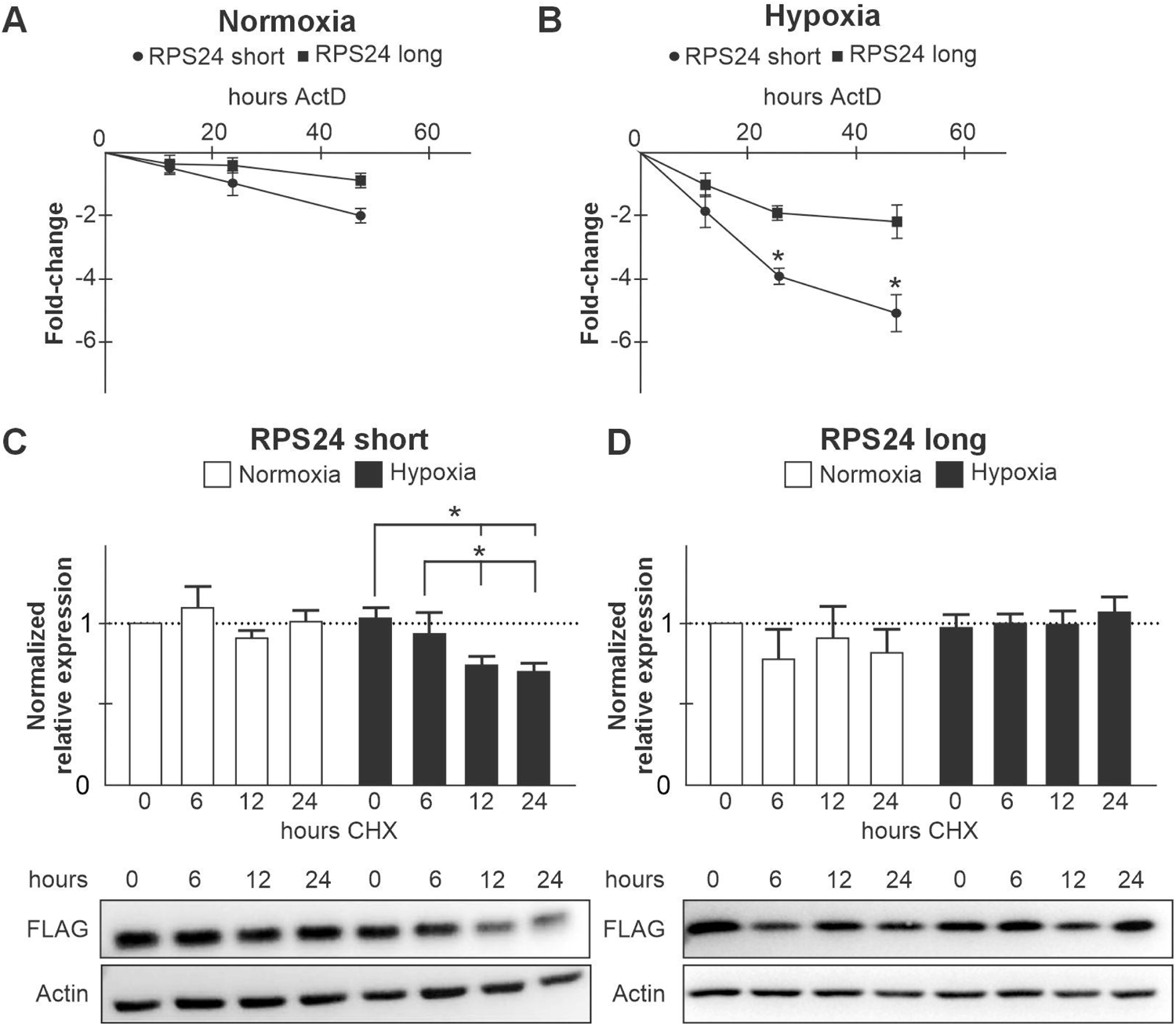
RPS24 long mRNA variant and the encoded protein isoform are more stable relative to the short form in hypoxia. RT-qPCR measuring the mRNA abundance RPS24 short (circles) and long (squares) transcript variants after 0, 12, 24, and 48 h of Actinomycin D (ActD) treatment in normoxic (A) and hypoxic (B) cells. The ΔΔCT method was used and the expression is represented as fold-change relative to 0 h. Cycloheximide (CHX) stability assay performed on (C) FLAG-RPS24 short and (D) FLAG-RPS24 long expressing cell lines cultured in normoxia or hypoxia for 0, 6, 12, 24 h. Representative western blot images of n=3. Densitometry calculated using Image Lab and normalized to β-actin loading control. Data (n ≥ 3) displayed as mean ± s.e.m. One-way ANOVA with Tukey’s HSD test was performed. * = P < 0.05. All experiments performed in U87MG glioblastoma cells.

Available antibodies against endogenous RPS24 cannot distinguish between the short and long isoforms due to the three amino acid difference. In a previous study, we generated stable cell lines expressing FLAG-tagged RPS24S and L protein isoforms to show that they could each associate with polysomes (13). Here, we performed a cycloheximide treatment to examine the stability of each RPS24 protein isoform in normoxia and hypoxia. The RPS24S protein isoform stability was unchanged during the assay in normoxia but was significantly decreased after 12 h and 24 h of hypoxia (Fig. 5C). The RPS24L protein isoform stability remained unchanged across the entirety of the assay in both normoxia and hypoxia (Fig. 5D). These data show that RPS24S mRNA, and the protein isoform it encodes, are less stable in hypoxia compared to RPS24L.

### Cell viability increases with exogenous RPS24L expression, but decreases with exogenous RPS24S expression in hypoxia

We next examined the impact of disrupting the RPS24L/S mRNA variant ratio on cell viability and growth in normoxia and hypoxia using Trypan Blue exclusion and crystal violet assays, respectively. As models of disrupted RPS24L/S ratios, we used the cell lines that stably express FLAG-RPS24L, FLAG-RPS24S, or empty vector control. Indeed, FLAG-RPS24S and FLAG-RPS24L expressing cells displayed a 30-fold lower and 40-fold higher RPS24L/S ratio, respectively, in both normoxia and hypoxia relative to control cells (Fig. 6A). Control cells displayed similar RPS24L/S ratios (Fig. 6A) to wild-type U87MG cells (Table 1) in normoxia and hypoxia. In normoxia, viability was constant among the three cell lines over the 72 h time course with the exception of FLAG-RPS24L overexpressing cells displaying lower baseline viability that was restored to 80% after 24 h (Fig. 6B). In hypoxia however, control cells displayed a significant decrease in viability from 80% to 50% after 24 h, which returned close to baseline levels at 72 h (Fig. 6C). FLAG-RPS24L expressing cells did not display a similar decrease in viability after 24 h, and in fact increased in viability keeping cells over 80% viable at 24 h and 72 h of hypoxic exposure (Fig. 6C). Conversely, FLAG-RPS24S expressing cells gradually decreased in viability to 50% at 72 h of hypoxia and did not recover like control cells (Fig. 6C).

**Figure 6.**
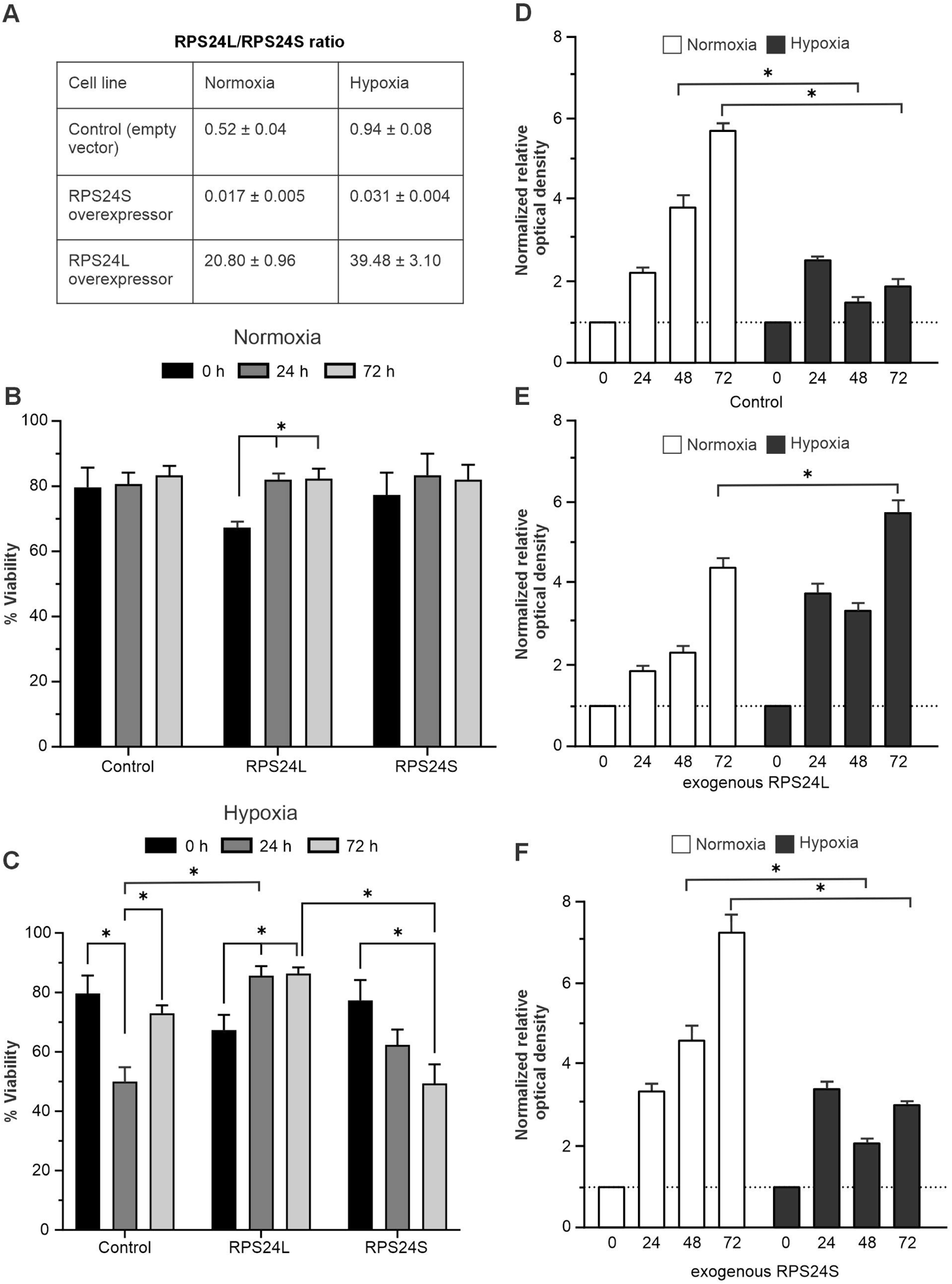
Cell viability increases with exogenous RPS24L expression, but decreases with exogenous RPS24S expression in hypoxia. (A) RPS24 long:short mRNA variant ratio measured by qRT-PCR in normoxic and hypoxic cell lines stably expressing FLAG-RPS24L, FLAG-RPS24S, and empty vector control. Trypan Blue exclusion was used to measure the percent viability in B) normoxic and C) hypoxic cells that stably express FLAG-RPS24S, FLAG-RPS24L, or empty control vector over a 24 h, 48 h, and 72 h. Crystal violet staining was used to measure cell number after 24 h, 48, and 72 h of normoxia or hypoxia in cells stably expressing (D) empty vector control, (E) FLAG-RPS24L, and (F) FLAG-RPS24S. Data (n ≥ 3) displayed as mean ± s.e.m. One-way ANOVA with Tukey’s HSD test was performed. * = P < 0.05. All experiments performed in U87MG glioblastoma cells.

With respect to cell growth, control cells doubled every 24 h in normoxia, but failed to grow after 48 h and 72 h of hypoxia (Fig. 6D). FLAG-RPS24L expressing cells grew more slowly relative to control cells in normoxia, but grew more rapidly in hypoxia including a doubling from 48 h to 72 h (Fig. 6E). FLAG-RPS24S expressing cells grew more rapidly in normoxia relative to control cells, but displayed similar stalled growth after 48 h and 72 h of hypoxia (Fig. 6F). These data suggest that overexpressing RPS24L aids cells to survive and grow in hypoxia, while overexpressing RPS24S hinders.

## DISCUSSION

Hypoxia-induced alternative splicing has been highlighted as a hallmark of cancer (14). Several studies have linked RPS24 to cancer progression but without considering splice variants. A single nucleotide polymorphism in the *RPS24* promoter was associated with risk of colorectal cancer (31). A genome wide association study identified RPS24 as a key gene within a survival network hub in uveal melanoma (32). Knockdown of RPS24 mRNA repressed cell migration and proliferation in colorectal carcinoma (33). A previous study from our group specifically linked the inclusion of a 22 bp cassette exon that produces the RPS24L mRNA variant to tumor hypoxia in prostate cancer (13). We also showed that the RPS24 protein isoform produced from this long mRNA variant can incorporate into actively translating ribosomes (13). Therefore, we set out to investigate how this alternative splicing event in RPS24 is induced and regulated, and how the protein isoform benefits cells.

Our data demonstrate that hypoxia, and the spheroid microenvironment, induces the RPS24L and represses the RPS24S mRNA variants. We show that these two variants are regulated by distinct pathways that both occur in hypoxia and spheroids: autophagy and HIF activity/mTOR repression. RPS24L hypoxic induction was repressed almost to baseline levels when autophagic flux was disrupted with Bafilomycin (Fig. 4D). Autophagy is an evolutionarily conserved process that modulates the recycling and reuse of energy and matter involving sequestration of cytoplasmic components within the autophagosome and subsequent delivery to lysosomes for degradation (34). Several stimuli can induce autophagy including hypoxia and acidosis, both features of the tumor microenvironment, to aid in normal and tumor cell survival (29,35). This is consistent with the data in Figure 2 demonstrating that low pH of 5.8 in normoxia and chronic hypoxia of seven days induced RPS24L independently. Further, serum starvation, a potent inducer of autophagy (28), induced RPS24L in normoxia by 1.5-fold (Fig. 4H). Importantly, serum starvation and hypoxia synergized to induce RPS24L by 2.52-fold (Fig. 4H), which was not observed when hypoxia and acidosis were combined, for example (Fig. 2A). Therefore, we propose that RPS24L is induced as part of a survival response when autophagy is activated. This would be consistent with the overexpression of RPS24L aiding (and RPS24S hindering) hypoxic cell survival (Fig. 6C) and previous studies linking RPS24 as a survival gene in uveal melanoma (32).

On the other hand, RPS24S repression was dependent on HIF stabilization and caused by mTORC1 inhibition, two phenomena that occur in hypoxia. While HIF stabilization and mTORC1 repression can be linked through the HIF-dependent transcription of the *REDD1* gene (3,36), we show that RPS24S repression can occur in both normoxia and hypoxia when HIFs are stabilized (Fig. 3A-C) or when mTORC1 is repressed (Fig. 4E and K) independent of one another. One could hypothesize that a HIF-dependent gene, such as a splicing factor, is directly or indirectly responsible for repressing the processing of RPS24S. Moreover, since many proteins of the small ribosomal subunit are involved in ribosome biogenesis (37), perhaps RPS24S is the favored isoform for this process and is repressed in hypoxia through energy stress-induced mTORC1 repression. Interestingly, even though mTORC1 inhibition induces autophagy, RPS24S repression through mTORC1 inhibition was not reversed upon the addition of Bafilomycin (Fig. 4E). This suggests that RPS24S mRNA is reduced under mTORC1 repression independent of autophagy, but that the hypoxic increase of RPS24L mRNA is dependent on autophagy but independent of mTORC1. The mTORC1-dependent induction of autophagy is well-characterized (27), but the mTORC1-independent induction of autophagy is less so. A study described an example of the latter where disruption of the endoplasmic reticulum-mitochondria calcium communication caused a bioenergetic crisis and mTORC1-independent AMPK-dependent autophagy (38). Therefore, one can appreciate that the parallel and distinct intracellular signalling cascades that induce RPS24L and repress RPS24S in hypoxia are fertile ground for further exploration.

We also probe the biological relevance of the hypoxic induction and repression of RPS24L and RPS24S, respectively. We show that the RPS24L mRNA and protein isoform are more stable than their RPS24S counterparts in hypoxia whereas they have similar stability in normoxia (Fig. 5). The RPS24L mRNA has an extra 22 bp cassette exon that contains predicted binding motifs (http://rbpdb.ccbr.utoronto.ca/) for several splicing factors, but also Smaug, which can recruit mRNAs to stress granules to protect them during stress (39). This 22 bp exon contains a premature stop codon generating a protein isoform three amino acids shorter than RPS24S, which contains an extra proline, lysine, and glutamate. Lysine is frequently targeted for post-translational modifications such as ubiquitin and acetylation that could influence protein stability and/or function. We show here that increasing the RPS24L/S ratio aids in hypoxic cell survival and growth (Fig. 6). Perhaps the more stable RPS24L aids ribosomes to maintain a level of protein synthesis required to mount a stress response. We have previously shown that both FLAG-RPS24L and FLAG-RPS24S can incorporate into translating ribosomes, presumably one or the other (13). Also, proximity proteomics of eIF4A1 showed enrichment of RPS24, highlighting its position near the mRNA entry channel and potential importance in translation initiation (40). Therefore, one could speculate that the RPS24 isoforms could be part of specialized ribosomes (12), an emerging concept, to direct ribosomes to certain transcripts as a response to stimuli.

Our data show that four cell lines from different tissues of origin display vastly different RPS24L/S baseline ratios (Table 1), but these ratios are increased in a similar manner in hypoxia and spheroids (Fig. 1). RPS24 is a ribosomal protein that displays tissue-specific alternative splicing with the long variant being most-predominant in the brain (the organ with the lowest mean partial pressure of oxygen (41)) and the short variant in liver and kidneys, for example (42–44). Tissues with a high RPS24L/S ratio could be better suited to respond to hypoxic stress or even stress in general. Cancer cells are notorious for resisting stress. We show that PC3 prostate cancer cells have a baseline RPS24L/S ratio of 3.6, by far the highest of the four cell lines tested (Table 1). Therefore, perhaps prostate cancer cells rely on a high RPS24L/S ratio to resist stress inputs. Indeed, prostate cancer cells have an innate resistance to fluid shear stress and the chemotherapeutic paclitaxel (45,46). Further, we have shown previously that RPS24L expression correlates with hypoxia in prostate cancer tissue (13).

These findings highlight that cells shift toward a long mRNA splice variant of RPS24 using parallel pathways that include autophagy, mTORC1 repression, and HIF stability to aid in survival. This has implications for a better understanding of basic cell biology, but also how cancer cells survive the tumor microenvironment.

## EXPERIMENTAL PROCEDURES

### Cell culture, reagents, and treatments

All cell lines were obtained from the American Type Culture Collection and maintained as suggested. These were tested for mycoplasma contamination and characterized by short tandem repeat and Q-band assays. Normoxic cells were maintained in a humidified chamber (ambient O_2_, 5% CO_2_, and 37 °C). Hypoxic cells were incubated for the indicated amount of time in a HypOxystation H35 (HypOxygen) at 1% O_2_, 5% CO_2_, and 37 °C. Hypoxic cells received pre-equilibrated (for at least 24 h) hypoxic media after being seeded 24 h prior. For long term cultures (up to seven days), cells were seeded at low density or from a serial dilution and the media replaced with fresh (pre-equilibrated for hypoxia) media every 48 h. Serum starved cells were cultured without fetal bovine serum or antibiotics for 24 h. Stable cell lines expressing FLAG-RPS24 isoforms and empty control vector were generated as previous described (13). Stable HIF constructs were obtained and transfected as previously described (47). The following drug concentrations were used for the indicated incubation time: Staurosporine (Abcam 146588) at 5 µM, DMOG (Cayman Chemical Co.) at 200 µM, Echinomycin (Tocris, NSC-13502) at 20 nM, TC-S 7009 (Tocris) at 100 μM, 10 mM Metformin and 50 mM of Oligomycin (kind gifts from Dr. Paul Spagnuolo, Guelph), and 200 μM CPI-613 (Cayman Chemical Co.), Torin at 1 µM (New England Biolabs), and Bafilomcyin at 10 nM (Sigma). Treatments on spheroids were done at 3-fold higher concentrations to the final 24 h of growth.

### Spheroid Formation and Treatment

Spheroids were generated by seeding 50,000 cells into round bottom, low-attachment 96-well plates (Corning) followed by gentle swirling to promote cell–cell contact and grown in normoxia for 1-7 days. Where specified, spheroids were treated by carefully removing media and replacing with drug-containing media for 24 h. For ethylene glycol tetratacetic acid (EGTA; Sigma E-3889) treatment, spheroids were incubated in normoxia at 37 °C for four days to allow proper formation. New media with or without 10 mM EGTA was added to the spheroids, which were incubated for an additional 24 h. Brightfield microscopy images were taken of the day 5 spheroids (Nikon Eclipse Ti Microscope). Spheroids were lysed for total RNA extraction, reverse transcription, and qPCR analysis of RPS24S and L mRNA abundance. For exposure of cell monolayers to spheroid media, five-day old spheroids were centrifuged at 4000 rpm for 90 sec after passing through a 27 gauge syringe. Trypan blue (Gibco) was used to confirm that the cells of the spheroids were not lysed. Cells were incubated with spheroid media for 3 h and 6 h before RNA extraction and qRT-PCR analysis.

### RNA Extraction and RT-qPCR

RNA was extracted from cell monolayers using Trizol (Invitrogen) per manufacturer’s instructions. For collection of RNA from spheroids, approximately 35 spheroids were collected and pelleted by centrifugation followed by washing with Phosphate Buffered Saline (PBS). Pelleted spheroids were resuspended in Trizol and passed through a 27-gauge needle. Reverse transcription of 2 µg RNA was completed using the High-Capacity cDNA Reverse Transcription Kit (Applied Biosystems) as per manufacturer’s instructions. Quantitative PCR was performed using SsoAdvanced Unviersal SYBR Green Supermix (BioRad) in a CFX96 PCR detection system using CFX manager software (BioRad). Relative fold change of expression was calculated using the ΔΔCT method with reference genes RPL13A and/or RPLP0 and normalized as specified. Primers: RPS24S Forward 5’-GACTTGCAAGACATGGCCTGT-3’, Reverse (5’-TCCTTCGGCTTTTTGCCA-3’; RPS24L Forward 5’-GACTTGCAAGACATGGCCTGT-3’, Reverse 5’-TCCAGCTCATTTTTGCCAG-3’; RPL13A Forward 5’-AGCCGCATCTTCTGGCGGA-3’, Reverse 5’-TTGTCGTAGGGCGGTGGGATG-3’; RPLP0 Forward 5’-AACATCTCCCCCTTCTCC-3’, Reverse 5’-CCAGGAAGCGAGAATGC-3’; GLUT1 Forward 5’-CTGTCTGGCATCAACGCTGTCTT-3’, Reverse 5’-TCCTCGGGTGTCTTGTCACTTTGG-3’; TGF-β2 Forward 5’-CACCATAAAGACAGGAACCTG-3’, Reverse 5’-GGAGGTGCCATCAATACCTGC-3’.

### Acidosis and lactic acid treatment

Cells were treated with acidosis-permissive media that is at neutral pH when added to cells, but decreasing buffer concentrations allow for a gradual decrease in media pH resulting from cellular metabolic processes (modified from (48)). Over 24 h, this resulted in a pH gradient where the media self-adjusted to a final pH as follows: 5.8, 6.0, 6.3, 6.5, 6.8, and 7.0. Powdered media was made with four concentrations of NaHCO_3_ (16 mM, 22 mM, 34 mM, and 40 mM) in two batches, one of which was adjusted to pH 6.0 (AP) and the other to pH 7.2 (SD). Air was bubbled into the pH 6.0 AP media until it increased to pH 7.2, as well as briefly bubbled into pH 7.2 SD media to stabilize the pH. The media were then sterilized by vacuuming through 0.2 µm bottle top filters. Serum (7.5% fetal bovine serum and 1% penicillin-streptomycin) was added to media following filtration. Use of the following media resulted in a pH gradient after 24 h of incubation with cells: 16 mM AP (pH 5.8), 34 mM AP (pH 6.0), 40 mM AP (pH 6.3), 22 mM SD (pH 6.5), 34 mM SD (pH 6.8), and 40 mM SD (pH 7.0). Cells (350,000) were seeded into 6-well plates 24 h before treatment, followed by subsequent addition of acidosis-permissive media and incubation in normoxia or hypoxia for 24 h before harvesting. For lactic acid treatment, cells were seeded and maintained in serum-free media for 24 h prior to treatment. Cells were treated for 24 h with 20 mM Lactic Acid pH 7.1 (Fisher Chemicals), 20 mM Sodium Lactate pH 7.4 (Fisher Chemicals) or 25 mM Sodium 57 Oxamate (Alfa Aesar).

### Transcript and protein Stability Assays

For transcript stability, cells were cultured as described in normoxia and hypoxia in the presence or absence of 10 μg/mL Actinomycin D (Sigma). Cells were lysed with Trizol at the specified time points for subsequent RNA extraction, reverse transcription, and qPCR to measure transcript variant abundance. For protein stability, FLAG-RPS24 L/S cells were cultured as described in normoxia or hypoxia in the presence or absence of 0.1 mg/mL cycloheximide (Fisher Chemicals). Cells were lysed with RIPA buffer (10 mM Tris-HCl pH 8, 1 mM EDTA, 0.5 mM EGTA, 1% Triton X-100, 0.1% sodium deoxycholate, 0.1% sodium dodecyl sulfate, 150 mM NaCl) at the specified time points for subsequent protein quantification and western blotting.

### Western Blotting

Standard western blotting procedures were used. Primary antibodies: HIF-1α (Novus Biologicals, NB100-123), HIF-2α (Novus Biologicals, NB100-122), LC3BP (Novus Biologicals NB100-2220), SQSTM1/p62 (Cell Signalling 5114), Flag (Sigma F1804), and β-Actin (GeneTex GT5512). HRP conjugated secondary antibodies anti-mouse (Promega, PR-W4012) and anti-rabbit (Invitrogen 65-6120). Densitometry was performed using Image Lab (BioRad).

### Crystal Violet Assay

FLAG-RPS24S, FLAG-RPS24L and empty vector control expressing stable cell lines were seeded into 12-well cell culture plates (Corning). After 24 h, cells were given fresh media and incubated for the indicated time. At each time point, cells were washed, fixed with methanol, and stained with 1% crystal violet solution in 20% methanol. The crystal violet solution was washed off and the plates were allowed to dry overnight. The next day cells were de-stained with 10% acetic acid and the absorbance of each well was measured at 595 nm using a ThermoMax microplate reader. This assay washes away dead cells before staining, and only the DNA and proteins of live adhered cells is stained.

### Trypan Blue Exclusion Assay

FLAG-RPS24S, FLAG-RPS24L and empty vector control expressing stable cell lines were seeded into 6-cm culture plates (Corning). After 24 h, cells were given fresh media and incubated for the indicated time. At the specified time points, the cells were scraped off into the media, pelleted by centrifugation at 4,000g, and resuspended in 200 µL of 1X PBS. The cells were stained with 0.4% Trypan Blue Solution (Gibco), placed on a haemocytometer, and both viable (unstained) and unviable (stained) cells were counted.

### Statistical Analyses

All statistical analyses were performed using GraphPad Prism 9.2. Statistical differences between treatments were evaluated by one-way ANOVA followed by Tukey’s HSD test.

### Supporting information

This article contains supporting information.

## Acknowledgments

We thank Drs. Stephen Lee (Miami) and Paul Spagnuolo (Guelph) for reagents. A.B. was supported by a Natural Sciences and Engineering Council of Canada Alexander Graham Bell Canada Graduate Scholarship. M.M. was supported by the Dr. Anne Innis Dagg Summer Research Scholarship and an Ontario Graduate Scholarship. This work was funded by the Natural Sciences and Engineering Council of Canada grant number 04807 to J.U.

