## Supplementary figures and images for "Autophagy-dependent alternative splicing event produces a more stable ribosomal protein S24 isoform that aids in hypoxic cell survival"

### Supplemental Figure 1

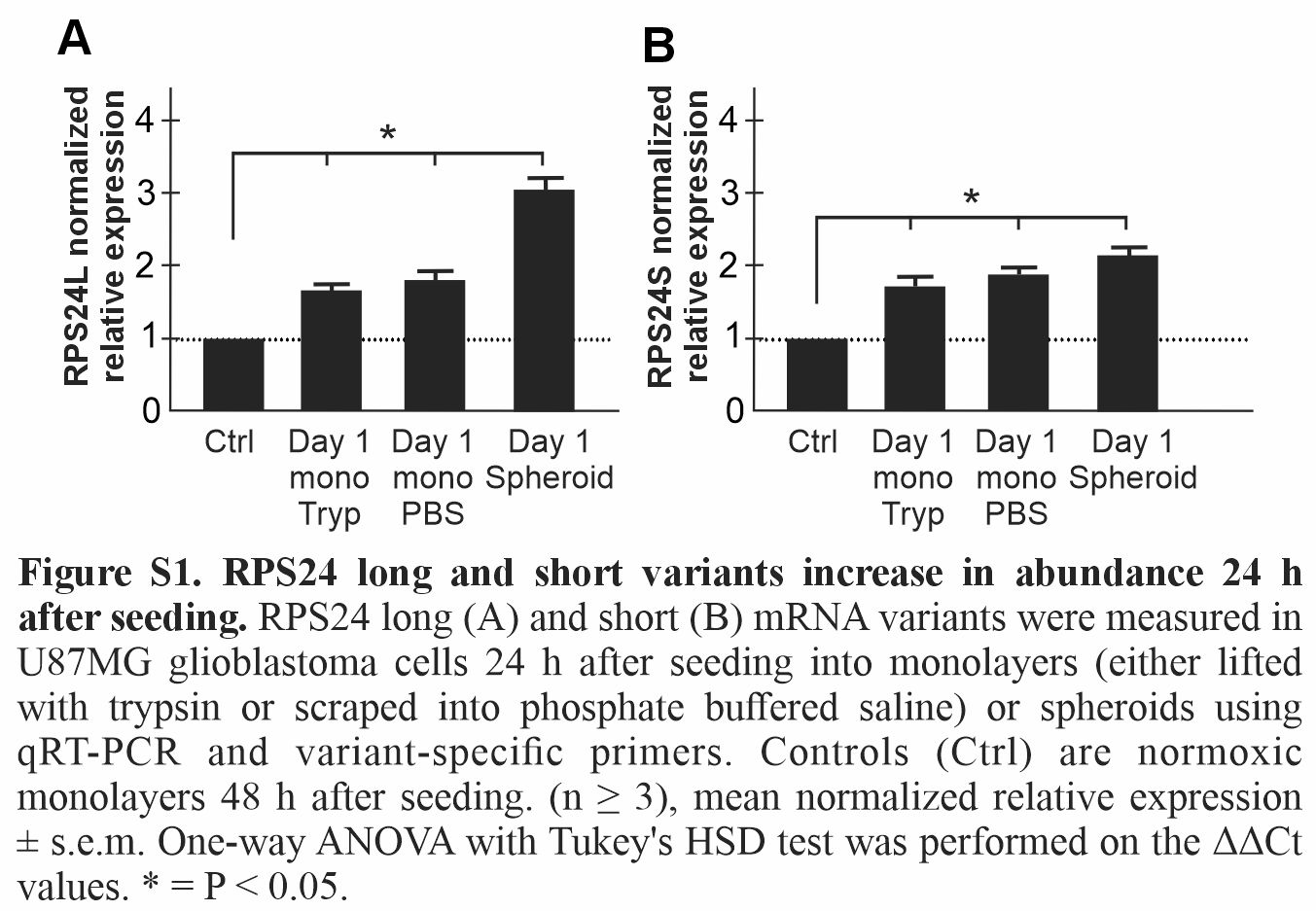
